# Accumulation of deleterious mutations during bacterial range expansions

**DOI:** 10.1101/093658

**Authors:** Lars Bosshard, Isabelle Dupanloup, Olivier Tenaillon, Rémy Bruggmann, Martin Ackermann, Stephan Peischl, Laurent Excoffier

**Affiliations:** CMPG, Institute of Ecology an Evolution, University of Berne, 3012 Berne, Switzerland; Swiss Institute of Bioinformatics, 1015 Lausanne, Switzerland; Interfaculty Bioinformatics Unit, University of Berne, 3012 Berne, Switzerland; INSERM, IAME, UMR 1137, Paris, France; Université Paris Diderot, Sorbonne Paris Cité, Paris, France; Institute of Biogeochemistry and Pollutant Dynamics, Swiss Federal Institute of Technology Zurich (ETH Zürich), 8092 Zürich, Switzerland; Department of Environmental Microbiology, Swiss Federal Institute of Aquatic Science and Technology (Eawag), 8600 Dübendorf, Switzerland

**Keywords:** Range expansions, mutation load, experimental evolution

## Abstract

Recent theory predicts that the fitness of pioneer populations can decline when species expand their range, due to high rates of genetic drift on wave fronts making selection less efficient at purging deleterious variants. To test these predictions, we studied the fate of mutator bacteria expanding their range for 1650 generations on agar plates. In agreement with theory, we find that growth abilities of strains with a high mutation rate (HMR lines) decreased significantly over time, unlike strains with a lower mutation rate (LMR lines) that present 3-4 times fewer mutations. Estimation of the distribution of fitness effect (DFE) under a spatially explicit model reveals a mean negative effect for new mutations (-0.38%), but it suggests that both advantageous and deleterious mutations have accumulated during the experiment. Furthermore, we show that the fitness of HMR lines measured in different environments has decreased relative to the ancestor strain, whereas that of LMR lines remained unchanged. Our results thus suggest that successful expanding species are affected by deleterious mutations that accumulate during the expansion process, leading to a drastic impairment of their evolutionary potential.

## Introduction

Beneficial mutations are generally viewed as the main driver of evolution through adaptation, but most species harbor many deleterious mutations that have surprisingly not been eliminated by selection (Agrawal and Whitlock 2012). These deleterious mutations are known to affect the rate of adaptation in asexuals (Denamur and Matic 2006; Lynch 2010; Orr 2000), shape patterns of neutral genetic diversity ((Charlesworth *et al.* 1995; Corbett-Detig *et al.* 2015), affect the evolution of recombination ((Gordo and Campos 2008; Keightley and Otto 2006) and mutation rates (Lynch 2010; Sung *et al.* 2012), and can lead to the extinction of sexual or asexual populations (Haigh 1978; Lynch *et al.* 1993). Nevertheless, the effects of deleterious mutations are commonly ignored in studies of adaptation (Fogle *et al.* 2008; Good *et al.* 2012; Weissman and Hallatschek 2014; Wilke 2004), probably because deleterious mutations are expected to contribute little to the evolutionary dynamics of large populations (e.g., Blundell *et al.* (2015), but see Covert *et al.* (2013)).

Most individuals harbor deleterious mutations in their genome (Fu *et al.* 2014; Garcia-Alonso *et al.* 2014; Henn *et al.* 2015; Tennessen *et al.* 2012; Xue *et al.* 2012), making them incur a mutation load (Kimura *et al.* 1963). While a high mutation load is expected in small populations (Kimura *et al.* 1963; Lynch *et al.* 1993), deleterious mutations are not necessarily restricted to low frequencies in large recombining populations. In humans, for instance, recent genome sequencing studies have shown a surprisingly high number of deleterious mutations, including loss of function mutations (Sulem *et al.* 2015). The exact processes responsible for the creation and preservation of a mutation load in demographically successful organisms with large population sizes are still unresolved, and remain a hotly debated subject in human population genetics (Do *et al.* 2015; Lohmueller *et al.* 2008; Simons *et al.* 2014). A central theme in this controversy is the effect of past demographic processes on the efficacy of selection and on current patterns of mutation load (Henn *et al.* 2015; Lohmueller 2014).

Theoretical studies have recently proposed that spatially expanding populations should accumulate deleterious mutations, due to small effective size and inefficient selection on the range margins, a phenomenon that was called expansion load (Peischl *et al.* 2013). The fitness of individuals on the front of the expansion is predicted to decrease over time and space (Peischl *et al.* 2013; Peischl and Excoffier 2015), potentially affecting the speed of the expansion and imposing constraints on the limits of a species range (Peischl *et al.* 2015). Intuitively speaking, repeated cycles of founder events and population growth occurring at the front of a range expansion lead to an evolutionary dynamic that is like that expected for populations undergoing recurrent bottlenecks. Range expansions are in some sense similar to mutation accumulation experiments, which attempt at removing the effect of selection by imposing strong and regular bottlenecks, often though a single individual (Trindade *et al.* 2010), but, unlike mutation accumulation experiments, range expansions are a continuous process where low densities on the front naturally limit the effect of selection.

However, whereas the process and consequences of expansion load during range expansions have been well described (Peischl *et al.* 2016; Peischl *et al.* 2013; Peischl and Excoffier 2015; Peischl *et al.* 2015; Sousa *et al.* 2014), direct empirical evidence for it is still lacking. Some theoretical predictions have however been supported by the analysis of human exomes. For instance, one observes that there is a clear increase of the number of sites homozygous for predicted deleterious mutations along the expansion axis out of Africa (Henn *et al.* 2016; Peischl *et al.* 2016), suggesting that the expansion process led to an increased recessive mutation load. Contrastingly, the additive load, as measured by the total number of derived alleles, seems rather constant in all human populations (Do *et al.* 2015; Simons and Sella 2016; Simons *et al.* 2014) (but see Fu *et al.* 2014), which is actually expected after range expansions (Peischl *et al.* 2016; Peischl and Excoffier 2015). Therefore, a more direct observation and clear measures of mutation load in successfully expanding species would thus be crucial to validate the theory.

Bacteria like *Escherichia coli* are an ideal candidate to directly test the prediction that expanding populations incur an expansion load. Indeed, *E. coli* growing on agar plates form sectors of low diversity where single mutants have fixed, which has been attributed to high rates of drift and low effective sizes on expanding wave fronts (Hallatschek *et al.* 2007; Korolev *et al.* 2010). Their mode of expansion on plate without much lateral movement should thus lead to strong differences between lines from different sectors (Peischl *et al.* 2016), and look like one dimensional expansions for which theoretical predictions have been made. Also, their haploid nature and asexual mode of reproduction simplifies the estimation of load, as no assumptions on the dominant-recessive status of harmful mutations are necessary, unlike in diploids (Simons and Sella 2016). Another advantage of *E. coli* is that strains with a defective mismatch repair system have a very high mutation rate (Barrick *et al.* 2009; Lee *et al.* 2012; Tenaillon *et al.* 2016; Trindade *et al.* 2010), increasing diversity and our ability to observe evolution of bacterial growth and reproduction over a relatively short time. One would thus expect to be able to see if a prolonged period of bacterial expansions leads to a decrease in population fitness over time and space as predicted by theory (Peischl *et al.* 2013; Peischl *et al.* 2015).

However, it is still unclear if rare beneficial variants could compensate the negative effects of more frequent deleterious mutations (Hallatschek and Nelson 2010; Lehe *et al.* 2012). Indeed rare positively selected mutations can increase in frequency on an expanding front more quickly than they would in a well-mixed population (Gralka *et al.* 2016b), and their recurrent fixation could potentially increase the fitness of front populations. Thus, evidencing the presence of an expansion load in naturally growing bacteria would oblige us to seriously reconsider the role that deleterious mutations play in evolution and adaptation of living organisms.

## Materials and Methods

### Range Expansion Experiment

#### Bacterial strains

We used *E. coli* K12 MG 1655 strains where the expression of the *mutS*gene is directly controlled by the arabinose promoter *pBAD* inserted in front of the *mutS* gene. In absence of arabinose, *mutS* is not expressed, leading to a higher spontaneous mutation rate due to the inactivation of the methyl-directed mismatch repair system (MMR, (Yang 2000)). Bacteria grown in presence of arabinose express the *mutS* gene and thus have a lower spontaneous mutation rate. Additionally, our strain had a GFP marker located in the *lac* operon, which can be induced by IPTG (Isopropyl β-D-1-thiogalactopyranoside) (see **Supplementary Fig. 1**).

#### Growth on agar plates

The growth of the mutator strain on agar plates was examined to find the time period during which the colony is expanding with a constant velocity on an agar plate. The mutator strain was grown overnight in liquid culture at 37°C in LB medium supplemented with 0.2% arabinose. One million cells were then deposited on five LB agar plates and incubated at 37°C up to 7 days and the colony size was measured every day. After a short period of exponential growth, the colonies were expanding with at a constant velocity on an agar plate for up to four days, and then decreasing.

All strains were grown on LB agar plates at 37°C for a total duration of 39 days. More precisely, we transferred strains on a new plate every three days, thus before their growth rate would begin to decrease (Fig. 1). An image of the colony was taken before transferring the cells to a new plate for later growth analyses. For each transfer, approximately 100 million bacteria were sampled from the front using a sterile pipette tip and re-suspended in 100 μl dilution solution (0.85% NaCl). The size of the sampling point to get ∼100 million bacteria was determined after an initial calibration. The number of sampled bacteria for different sampling sizes was first determined by plating serial dilutions on LB agar plates and incubation for 24 h at 37°C. The number of colony forming units (CFU) were counted on plates where the range of CFU was between 30 and 300. The CFUs were divided by the dilution factor to determine the number of bacteria present in the original solution. The location of the sampling point of each transfer was chosen at random on the periphery of the colony. New plates were then inoculated using 1 μl of this solution corresponding to ∼1 million cells. Note that this large initial population size guarantees that there is no bottleneck induced by the transfer itself, unlike what happens during mutation accumulation experiments, where bottlenecks are induced through a single clone. Selection, if any, should thus not be relaxed during this transfer, except for extremely mild mutations with selection coefficient smaller than 1/*N* or about 10^-6^. Then, 43 μl glycerol (50%) was added to the remaining bacteria suspension and bacteria were stored at −80°C. This range expansion experiment was done twice independently. The first experiment included the expansion of 48 high mutation rate (HMR) strains. The second experiment included the expansion of 9 HMR strains, and of 10 strains with a low mutation rate (LMR). In those LMR strains the MMR mechanism was only partially induced by adding 0.2% arabinose. The HMR lines from the first and second experiments have then been analyzed jointly.

**Figure 1:**
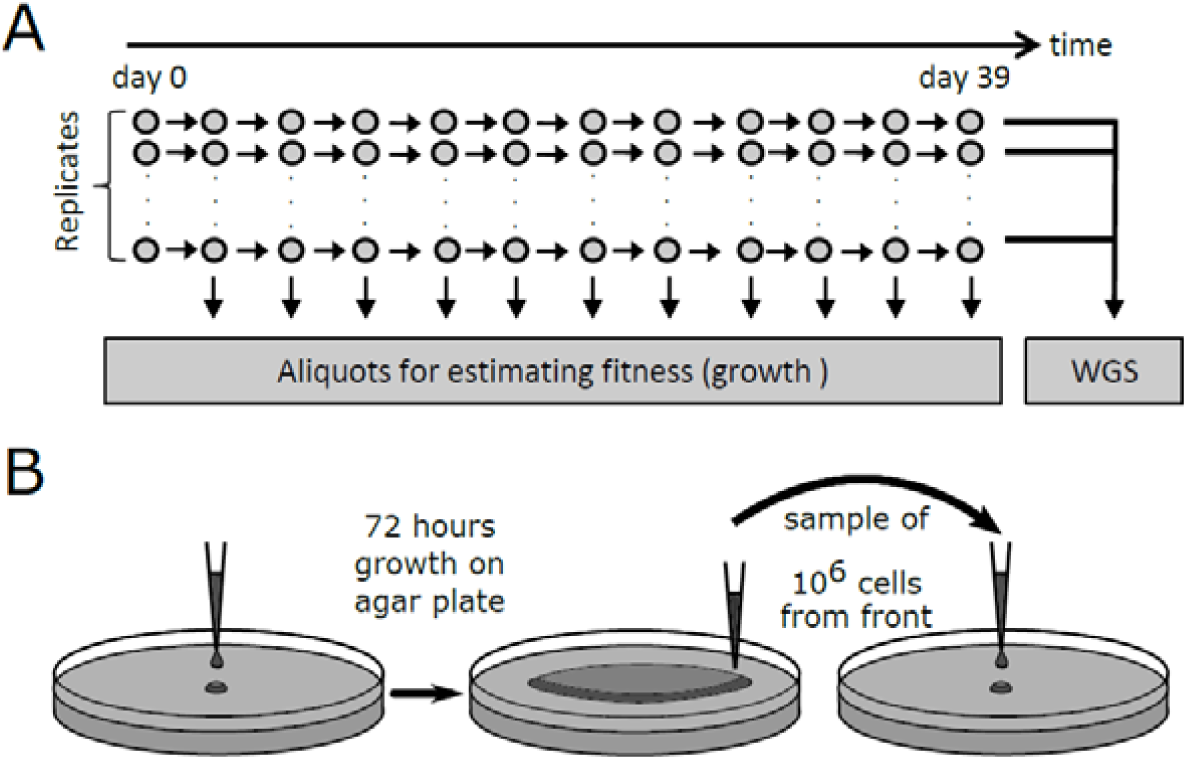
Experimental setup. **A:** *mutS-E. coli* lines were grown on agar plates for a total of 39 days or about 1650 generation assuming a generation time of 34’ (**Supplementary Fig. 2**). **B**: After 3 days of growth, about ∼100 million bacteria are sampled on the edge of the colony, diluted in 1 μl LB medium, and ∼10^6^ bacteria are deposited at the center of a new agar plate for a new 3-days growth cycle. This transfer should thus not impose any substantial bottleneck for the cells sampled on the edge of the colony that we are interested in following through time. This procedure was repeated 12 times for a total of 39 days of evolution for each line.

#### Generation time on the wave front

The generation time of bacteria at the front of the expanding colony was determined by using a confocal microscope (Leica TCS SP5) (**Supplementary Fig. 1**). Cells from −80°C glycerol stock were grown in LB medium at 37°C for 24h. 1 μl of this culture was transferred to an LB agar plate containing 0.1 mM IPTG and incubated at 37°C for 24h. A picture of the front of the colony was taken every two minutes for one hour with a 63x dry objective. The temperature during the measurement was set to 37°C by using an incubation chamber. The images were analyzed by using the Fiji (Schindelin *et al.* 2012) image analysis software. The cell mass increase is directly proportional to the length of the cell (Kiviet *et al.* 2014), which was measured by analysis of fluorescence intensity profiles along the cell axis. This experiment was done three times independently. In total the length increase of 49 individual cells was measured. The growth rate was determined by calculating the elongation rate by using a linear mixed-effect model *L* = *L*_0_ × 2^*rt*^, where *L* is the length of the cell, *L*_0_, is the length of the cell at time 0, *t* is the time in minutes, and *r* is the growth rate. The elongation dynamics of 16 cells is shown in **Supplementary Fig 2**. The average generation time corresponding to a doubling in cell mass estimated from the mixed-effect regression analysis is 34.2′, 95%CI [33.0, 35.6]. This generation time was used to estimate mutation rates per generation

### Estimation of mutation rate with a fluctuation test

A fluctuation assay (Foster 2006) was used to calculate the mutation rate of the ancestral strain, three HMR strains and one LMR strain. The rate at which mutations occur to enable cells to grow on selective agar was calculated. For each strain, we used 45 independent cultures. The ancestor strain and the HMR strains were incubated in LB medium without arabinose and the LMR strain in LB medium with 0.2% arabinose. The starting concentration of the cultures was 5 10^4^ cfu/ml, and the end concentration of the cultures was 1.5 10^9^ cfu/ml. The bacteria (initially not resistant to nalidixic acid were then exposed to LB agar plates containing 100 μg/ml nalidixic acid. Resistant colonies were counted and the data was analyzed with the MSS-Maximum Likelihood Estimator Method using FALCOR (Hall *et al.* 2009).

### DNA sequence analyses

#### DNA extraction

After the range expansion experiment on agar, one million cells from the wave front were streaked out on an LB agar plate containing 0.2% arabinose and incubated for 24h at 37°C to isolate single clones. A single colony was dissolved in 100 μl dilution solution (0.85% NaCl) and 1 μl was transferred to a new LB agar plate containing 0.2% arabinose. The plate was then incubated for 24h at 37°C. Then, the entire colony was removed from the agar plate and resuspended in 1 ml dilution solution. Genomic DNA was extracted using the Wizard Genomic DNA Purification Kit (Promega) following the manufacturer protocol. The integrity of the DNA was checked by gel electrophoresis. The DNA concentration was determined by fluorometric quantification (Qubit 2.0).

#### Whole genome sequencing

We sequenced DNA samples in two separate runs. 48 HMR samples were first sequenced using a TruSeq DNA PCR-Free library (Illumina) on a HiSeq2500 platform (Illumina), from which we obtained 100bp end reads for all samples. 9 HMR strains and 10 LMR strains were then sequenced using a paired end NexteraXT DNA library (Illumina) on a MiSeq platform (Illumina). The MiSeq platform generated 300bp end reads. Note that we did not find any differences between the average number of mutations (*m̅*) for HMR lines sequenced on the HiSeq2500 or MiSeq platforms (*m̅* = 113.8.4 vs. 121.4, respectively, t-test, p=0.247)

#### Neighborjoining tree

We computed genetic distances between each pair of samples using point substitutions and the Kimura 2-Parameters distance metric (Kimura 1980). The distance matrix was then used to build a neighbor joining tree with Phylip version 3.695 (Felsenstein 2005), which is represented in Fig 2.

**Figure 2:**
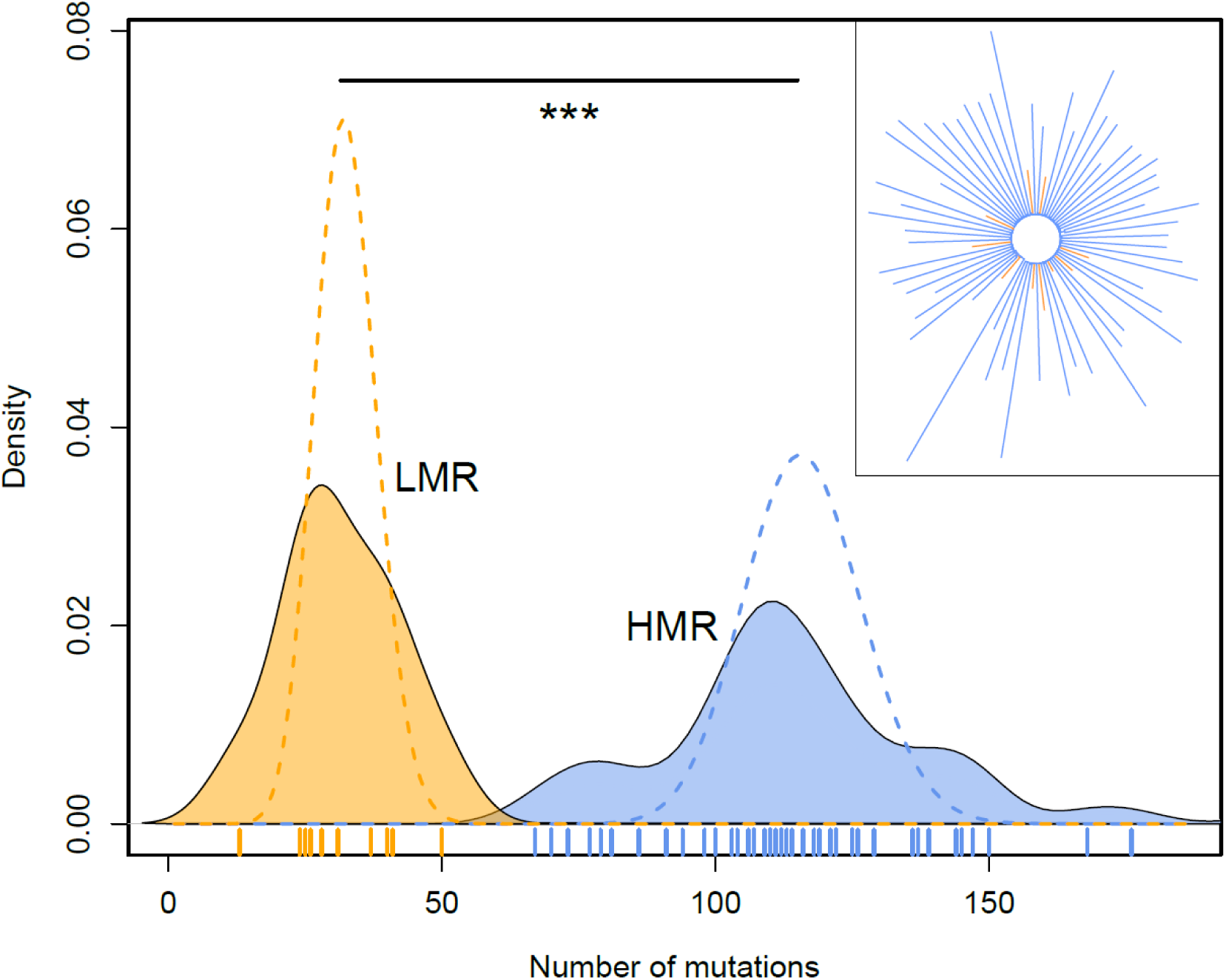
Number of mutations in evolved strains. Distribution of the total number of observed mutations per strain. Dashed lines are Poisson distributions fitted to the mean of the observed distributions. The two means are significantly different by a Mann-Whitney test (p-value =5.52×10^-7^). In the upper right inset we show a neighbor joining tree of the different strains (represented with the same color code as in the main figure).

#### Variant calling

We used Trimmomatic 0.32 (Bolger *et al.* 2014) to remove the adapter sequences from the reads and for quality trimming. Leading and trailing bases with quality below 3 were removed. The reads were scanned with a 4bp sliding window, and cut if the average quality per base was below 15. Reads with a length below 36 were excluded from the analysis. Burrows-Wheeler Aligner (BWA) 0.7.5 (Li and Durbin 2009) was used to map the reads to the *E. coli*K12 MG 1655 reference genome (NCBI Reference Sequence: NC_000913.3). Picard tool 1.99 was used to remove PCR duplicates and variants calling was performed with Genome Analysis Toolkit (GATK) 2.7 (McKenna *et al.* 2010). The SNPs were filtered based on the following VCF field thresholds: QD < 2.0, FS > 60, MQ < 40. Indels were filtered based on the following VCF field thresholds: QD < 2.0, FS > 200. Substitutions and small indels were also called using a modified version of a previously published pipeline (Tenaillon *et al.* 2012). We only kept mutations if the proportion of reads carrying the variant was larger than 75%. Substitutions and indels were retained as independent events if they could not be attributed to a gene conversion event. We used as a signal of gene conversion the presence of the mutated sequence (the mutated base and its 30bp neighboring bases) somewhere else in the genome. Mutations within 200 bp of a gene conversion signal were also considered as gene conversion mutations. All mutations were manually validated thanks to a visual output and kept for further analysis if they were both detected with GATK and the pipeline described above. SnpEff 4.0 was used to annotate the variants (Cingolani *et al.* 2012; Foster *et al.* 2013).

#### Estimation of *dN/dS* ratio

We computed the ratio of non-synonymous over synonymous substitutions (*dN/dS*) for the bacterial lines by counting the number of non-synonymous and synonymous substitutions accumulated in each line as compared to the reference sequence, as well as the number of non-synonymous and synonymous substitutions expected if all codon positions in the reference sequence would mutate. *dN/dS* can then be written as *dN* / *dS* = ∑*N_obs_* / ∑(*S_obs_N_tot_* / *S_tot_*), where *N_obs_* and *S_obs_* are the numbers of observed non-synonymous and synonymous substitutions in a given line, and *N_tot_* and *S_tot_* are the total number of expected non-synonymous and synonymous substitutions, respectively, over the whole bacterial genome. The observed and expected numbers of non-synonymous and synonymous substitutions in each line were computed for each 4 possible transitions and 2 possible transversions and summed across the 6 categories of mutations. We used a bootstrap approach to compute *dN/dS* confidence intervals for each strain. Briefly, *dN/dS* was computed with the above-mentioned approach using randomized datasets in which the mutations were randomly sampled with repetition among the 12 observed categories of mutations (6 types of synonymous and 6 types of non-synonymous mutations). To test for differences between treatments, we used a permutation scheme to obtain the null distribution of the amount of differences in average *dN/dS* values between treatments, taking sample size differences into account.

#### PROVEAN scores

We used PROVEAN scores (Choi and Chan 2015; Choi *et al.* 2012) to compute the potential damaging effects of non-synonymous substitutions and in-frame insertions and deletions observed in bacterial lines. PROVEAN scores are alignment-based scores measuring the change in protein sequence similarity before and after the introduction of the amino acid variation to the mutated sequence (Choi and Chan 2015; Choi *et al.* 2012). A threshold of −2.5 for PROVEAN scores has been shown to allow for the best separation between deleterious and neutral classes of variants in human and non-human UniProt protein variation (Choi *et al.* 2012). PROVEAN scores below −2.5 are thus indicative of a severe and thus potentially deleterious effect of the mutation (Choi and Chan 2015; Choi *et al.* 2012), even though we cannot dismiss the possibility that they might be advantageous in some lines. Conservatively, we will assume that mutations with PROVEAN score below −2.5 have a strong phenotypic effect. The distribution of PROVEAN scores for HMR and LMR lines are shown in **Supplementary Fig 3**.

### Measures of fitness

#### Expansion velocity on agar plate

Images of the colony were taken during the experiments on agar plates (n=57 for HMR lines, n=10 for LMR lines) before transferring the cells to a new plate. We took a picture every three days for each line, and thus have a total of 13 pictures for each line. The images were analyzed with the Fiji package of the imageJ software (Schindelin *et al.* 2012). The radius of the colony was measured and plotted against time (Fig. 3). Note that points for day 12 in HMR lines from the first experiment were not considered in further analyses due to a potential batch effect. The change in expansion velocity was then determined by fitting a mixed-effect linear model to the data. This model assumes that the growth rate of all lines changes due to a fixed effect common to all lines, but it considers line-specific variability in growth rates. The slopes of the regression lines plotted in Fig. 3 for each line are obtained by adding the fixed and line-specific effects.

**Figure 3:**
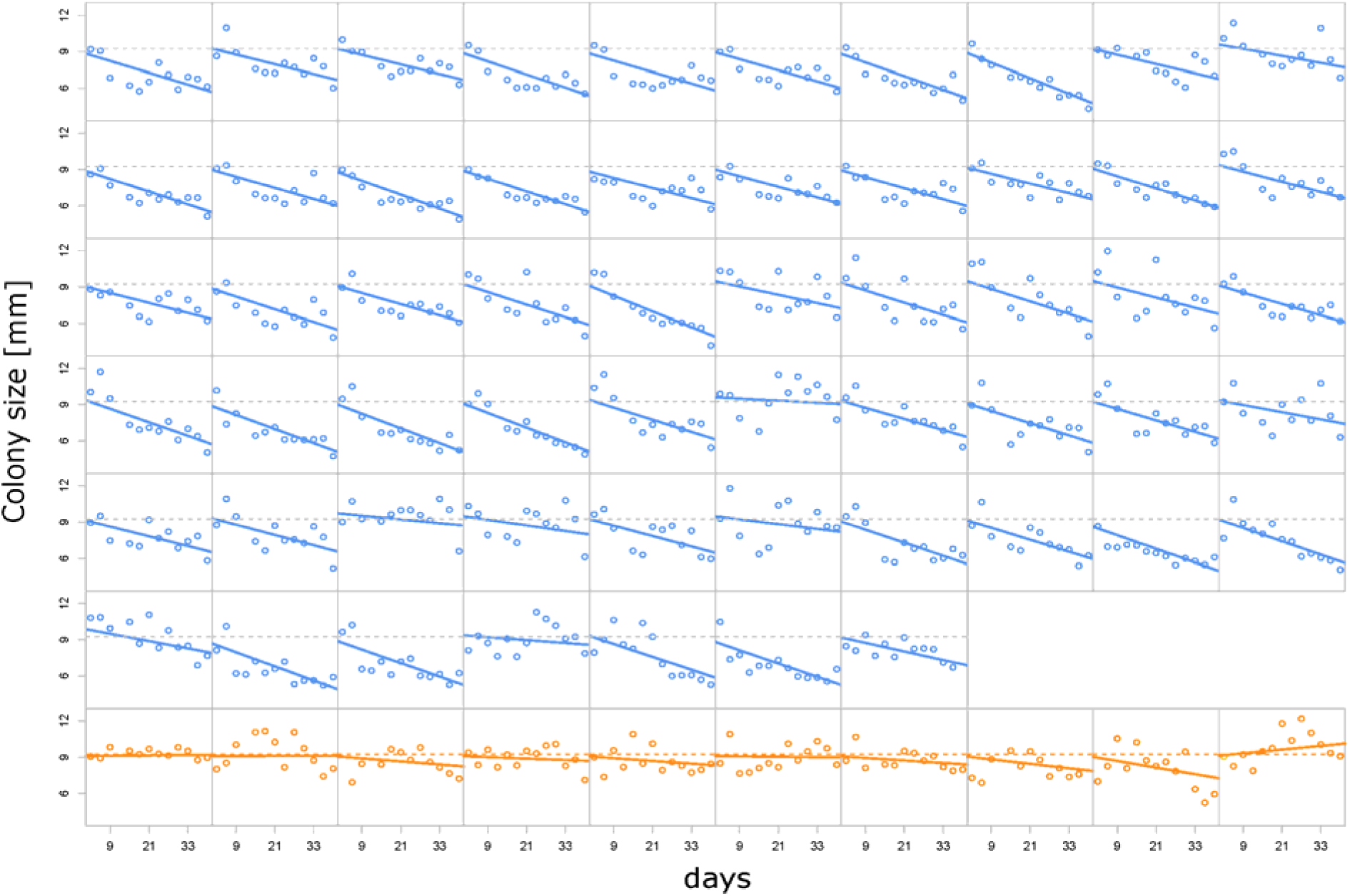
Evolution of colony radius after 3 days of growth on agar. Blue: HMR lines. Orange: LMR lines. The x-axis scale represents total days of evolution. Horizontal dashed lines represent the average colony size measured after the first 3 days of growth over all HMR or all LMR lines. Mixed effect linear regressions have been performed separately for HMR and LMR strains. Solid lines represent strain-specific regression lines, with slopes obtained as the sum of fixed and line-specific effects. HMR change in growth rate per day: −78 μm, 95%CI [−85; −70], p-value<2e-16. LMR change in growth rate per day: −11 μm, 95%CI [−33; 10], p-value=0.29 NS.

#### Competition experiment on agar plates

##### Linear growth of unmixed ancestral and evolved strains.

To determine the change of fitness on agar plate, we growth-competed our evolved lines and their ancestral strain against a reference strain where the *lacZ* was deleted. The evolved strains could thus be distinguished from the reference strain by adding *x-gal* to the cells. Bacterial with a functional *lacZ* gene turned blue within 15 min whereas the reference strain stayed white (see **Supplementary Fig. 4**). An LB agar plate with 0.5% arabinose was inoculated with an evolved strain and with the reference strain. The cells were deposited linearly on the agar plate with a razor blade dipped in the bacterial culture, with the reference strain being placed next to the evolved strain, without mixing. The plates were incubated at 37°C for three days. After the growth 0.1M x-gal solution was sprayed on top of the agar plate and the plate was incubated for 15 min at 37°C. Images were taken and analyzed with Fiji (Schindelin *et al.* 2012). Capitalizing on the fact that it takes the same amount of time for the less fit strain to grow along the expansion direction as it takes the fitter strain to grow a longer distance, the relative fitness can be estimated by measuring the angle between the two strains (Korolev *et al.* 2012), as

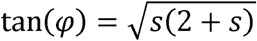

where *s* is the selection coefficient associated with the evolved strain, and □ is the angle between the two strains (see **Supplementary Fig. 4**). The selection coefficients of all evolved strains were then normalized by the mean selection coefficient of the ancestral strain, which was also competed against the reference strain. The results are shown in Fig. 4B.

**Figure 4:**
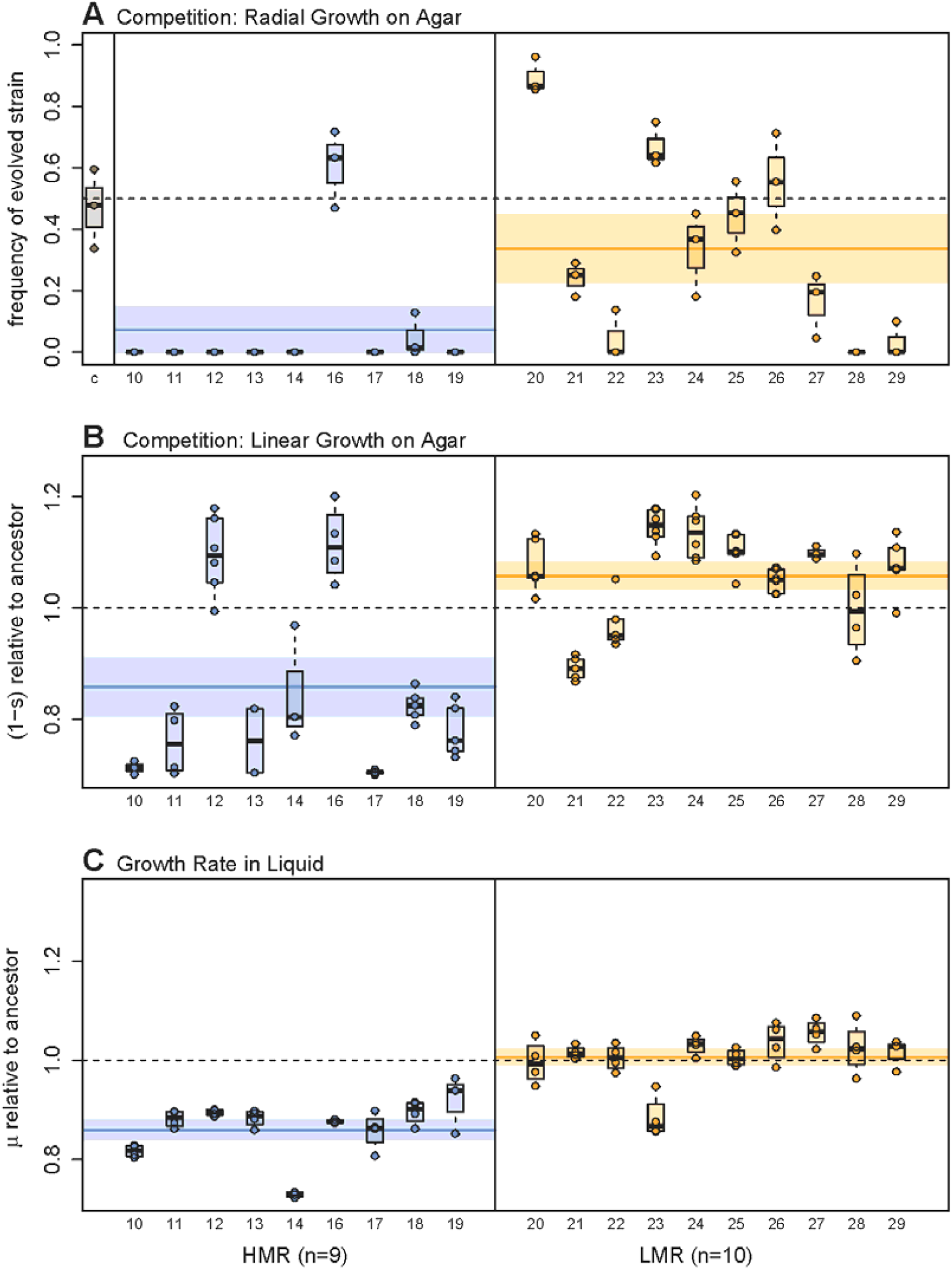
Estimation of bacterial fitness. **A:** Relative frequency of evolved strains on the edge of the colony after 3 days of radial growth on an agar plate, under conditions similar to those of our experiment of range expansion on agar (see Methods and **Supplementary Fig. 5**). Note that this measure only gives a qualitative assessment of the relative fitness of two strains, as this proportion will quickly change over time in case of unequal fitness (Gralka *et al.* 2016a). The first column (c) of the leftmost pane represents the relative frequency of the ancestral strain containing the same plasmid as the evolved strains, showing that the incorporated plasmids do not induce any fitness difference between strains. **B:** Competition on agar plate between a reference strain and evolved lines growing side by side for 3-days. The fitness of evolved strains (lines) is measured by the difference in growth rates at the contact zone between strains following Korolev et al. (2012). The angle formed by the contact zone between strains is indeed proportional to their fitness difference (see Methods). Each dot corresponds to one measure for a given strain. Note that the two HMR lines with highest fitness have a both a non-synonymous mutation in the mlc (makes large colonies) gene. **C:** Fitness of evolved strains relative to the ancestral strain, measured as growth rate in liquid culture. Note that labels on the x axis represent line ids.

##### Radial growth of well mixed ancestral and evolved strains

The ancestor strain was labeled with either GFP or mCherry containing plasmids, each plasmid having an additional ampicillin resistance gene. The strains from experiment 2 (9 HMR, 10 LMR evolved lines) were labeled with mCherry plasmid. All strains were grown in LB medium with 0.2 % arabinose and 50 μg/ml ampicillin at 37°C for 24 h. The density of the strains was adjusted by measuring the optical density and diluting the bacterial suspension with 0.85% NaCl solution to a final concentration of 10^9^ cfu /ml. The evolved strains were mixed with the GFP labeled ancestor strain in a 1:1 ratio. Additionally, we performed a control experiment where the mCherry labeled ancestor strain was mixed with the GFP labeled ancestor strain. 1 μl of each mixture was put in the center of an LB agar plate with 0.5 % arabinose and 50 μg/ml ampicillin and the plates were incubated at 37°C. After 3 days, a picture of the colony was taken and the fraction of the front occupied by the evolved strain was determined by using ImageJ, as shown in **Supplementary Fig 5**. This experiment was done three times independently for each sample. The results are reported in Fig. 4A.

#### Growth rate in liquid medium

Nine HMR and ten LMR lines were pre-grown in LB medium at 37°C until they reached an optical density of 0.2. The cells were diluted in fresh LB medium 1:100 and transferred to a 96-wells plate. A Spectrophotometer (BioTek Eon Microplate Spectrophotometer) was used to measure the log transformed optical density of the cultures every 30’ for 24h at 37°C. The exponential growth period was determined as the range where the logarithm of the optical density increased linearly (1h to 4h). The growth rate during this period was determined by computing the slope of a linear regression model. Four replicated measures were performed per line and reported in Fig. 4C.

#### Estimation of the distribution of fitness effects (DFE)

##### Estimating the DFE from bacterial growth dynamics

Peischl et al. (2015) developed an analytically tractable model for the evolution of mean fitness at an expansion wave front during a liner expansion along an array of discrete demes. One can use this model to predict the evolution of mean fitness and, assuming hard selection, also the evolution of colony size over time. Indeed, Peischl et al. (2015) showed that the change in mean relative fitness at the expansion front can be approximated using the following equation

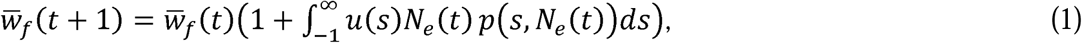

where *u*(*s*) is the mutation rate of mutations with effect *s*, *N_e_*(*t*) is the effective population size at the expansion front at time *t*, and *p*(*s*, *N_e_*) is the fixation probability of mutations with effect *s* at the front of an expanding population with an effective size of *N_e_*. The parameter *N_e_* measures genetic drift and corresponds to the compound parameter *FT* in Peischl et al. (2015), which is the product of the effective number of founders (*F*) and the time taken to fill an empty deme at the expansion front (*T*). Note that *T* is simply the inverse of the expansion speed. For definiteness and without loss of generality, we set the relative fitness at the onset of the expansion to *w̅_f_*(0) = 1. Note that *N_e_* is a parameter that depends on the expansion speed as well as the rate of migration of individuals, and hence also depends on mean fitness and may change over time (see Peischl et al. 2015 for details). Assuming exponential growth and that the growth rate is proportional to mean fitness one can show that in this model

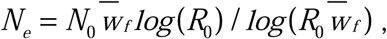

where *R*_0_ and *N*_0_ are the growth rate and effective population size at the expansion front at time *t* = 0. We have set *R*_0_ = 2, which simply defines the length of one generation as the time unit required for exponentially growing individuals to double their number.

We used a total mutation rate of 0.2 per individual per generation, as estimated by a fluctuation test (**Supplementary Table 4**). The distribution of fitness effects is conveniently modeled as a displaced gamma distribution (Shaw *et al.* 2002), as such a distribution naturally arises under Fisher’s geometric model (Fisher 1930; Martin and Lenormand 2006). In addition to the shape and scale parameters *α* and *β* of a conventional gamma distribution, this displaced distribution requires an additional parameter *δ* that represents the maximum effect of beneficial mutations, so that the distribution of individual mutation effects is given by

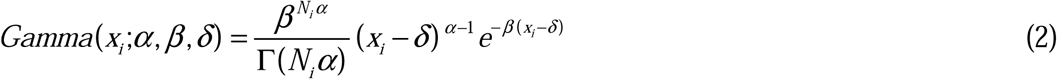

This leaves us with four unknown parameters to estimate: three parameters for the DFE and the effective population size at the expansion front at the beginning of the experiment, *N*_0_.

We then used the colony size trajectories (Fig. 3) to estimate the unknown parameters of this model. More precisely, we used the theoretical model to predict the expected evolutionary trajectory of the colony size by iterating eq. (1), assuming that the mean fitness at the expansion front is proportional to colony size. We then calculated the sum of squared deviations (SSD) between the 64 HMR lines and the theoretical expectation. A grid search over the parameter space was then used to determine the parameter combination that minimizes the total SSD of all 57 HMR lines. Note that we did not use the LMR lines here because the relatively small changes in colony size over time lead to numerical problems in the estimation procedure.

##### Estimating the DFE from the number of observed mutations

An alternative way to estimate the DFE is to use the relationship between the number of observed mutations and the fitness of the different lines. Assuming that mutation effects are additive, the total mutation effect *y_i_* of *N_i_* mutations that have accumulated in a bacterial strain and thus equal to 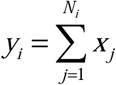 should also follow a displaced Gamma distribution similar to eq. (2), since the sum of Gamma variates also follows a Gamma distribution with rate parameter *β*, displacement parameter *δ*, but with a new shape parameter *N_i_α*. Therefore

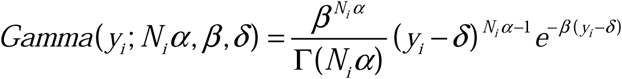

The likelihood *L*(*y*, **N**; *α*, *b*, *d*) of a set of *M* lines having accumulated **N** = {*N*_1_, *N*_2_, …, *N_i_*,…,*N_M_*} mutations is thus simply

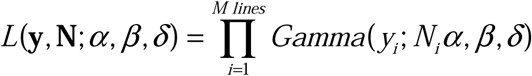

where **y** = {*y*_1_,…, *y_i_*,…,*y_M_*} is a vector of the sum of mutation effects estimated as one minus the fitness of the different lines. Taking the derivative of the log likelihood for *β* and equating it to zero allows us to get a maximum-likelihood estimator of *β* that only depends on *α* and *δ* as

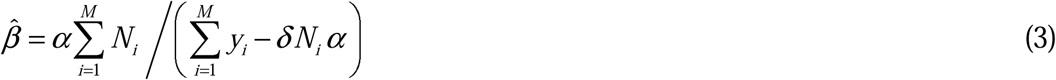

This ML estimator can be reinserted into the log likelihood equation, which becomes

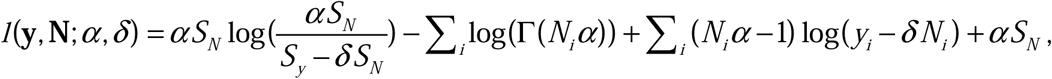

where *S_N_* = ∑_*i*_*N_i_* and *S_y_* = ∑_*i*_*y_i_*. The values of *α* and *δ* maximizing *l*(**y**, **N**; *α*, *δ*) can be found numerically, for instance by a simple grid search in the (*α*, *δ*) parameter space, from which eq. (3) can then be used to get the maximum likelihood estimate of *β*.

This procedure was first applied to the change in colony size of evolved lines between day 3 and day 39. In that case the total mutation effects *y_i_* was estimated as *y_i_* = 1 − *w_csi_*, where *w_csi_* = *CS*_3_ / *CS*_39_, and *CS*_3_ and *CS*_39_ are the colony sizes at day 3 and day 39 as predicted by a mixed effect regression model, respectively (see Fig. 3).

##### Testing for differences between HMR and LMR DFEs

We used a likelihood ratio test to test for potential differences between DFE’s separately estimated on HMR and LMR strains. If *l*(**y**, **N**; *α*, *δ*)_*All*_ stand for the log-likelihood computed on both HMR and LMR strains, and *l*(**y**, **N**;*α*, *δ*)_*HMR*_ and *l*(**y**, **N**;*α*, *δ*)_*LMR*_ stand for the likelihood of HMR and LMR strains, respectively, then the statistic *LogLR* = 2(*l*(**y**, **N**; *α*, *δ*)_*HMR*_ +*l*(**y**, **N**; *α*, *δ*)_*LMR*_ −*l*(**y**, **N**; *α*, *δ*)_*All*_) should be distributed as a *χ*^2^ with 3 degrees of freedom under the hypothesis that the DFEs for HMR and LMR are identical.

## Results

We used the bacterium *Escherichia coli* as a model system to test whether deleterious mutations can accumulate during the natural range expansion of an organism. We worked with a mutator (*mutS*-) strain in which we could experimentally modulate the mutation rates and thus analyze the evolutionary dynamics of mutation accumulation over a relatively short time (see Methods). We evolved replicated populations of *E. coli* for 39 days (about 1650 generations, **Supplementary Fig. 2**) by letting them expand radially across a solid substrate) for 13 periods of 3 days (Fig. 1 We performed this experiment with 57 lines having a high mutation rate (HMR lines) and with 10 lines having a lower mutation rate (LMR lines) (see Methods). Colony size of HMR and LMR lines was measured before each transfer and at the end of the 39 days, and aliquots of bacterial samples were preserved for whole genome sequencing (Fig. 1A).

### Genome-wide mutation patterns

We sequenced HMR and LMR lines at high coverage (>100X, **Supplementary Table 1**) to detect any potential differences in the number and patterns of mutations. After 39 days of range expansion, HMR lines accumulated ∼3.7 times more mutations than LMR strains (115.0 vs. 31.5 mutations per line, respectively, p<5.52 10^-7^, Mann-Whitney test, HMR range [67-204], LMR range [13-50], Fig. 2, **Supplementary Table 2**). When comparing substitutions between lines, we found that most of them (99.2%) occurred in distinct lineages and thus represent independent mutations (Fig. 2). Note that at first sight the mutation rate seems to have fluctuated along the genome (**Supplementary Fig. 6**), which appears mainly due to positional constraints during chromosome replication (**Supplementary Fig. 7**), suggesting that even though mutations are not randomly spatially distributed, they occurred randomly given local mutation rates constraints. Looking at the distributions of the number of mutations accumulated by each line, we observe that they are over-dispersed as compared to Poisson expectations (Fig. 2), suggesting that some mutator lines might have acquired genetic changes that further modify the mutation rate (Lee *et al.* 2012). We could confirm this hypothesis in the case of one HMR line containing no less than 176 substitutions and 17 frameshift mutations. In this line, we indeed identified a frameshift mutation in the *mutT* gene, whose inactivation specifically increases otherwise rare A:T->C:G transversions (here n=85) (Fowler and Schaaper 1997) that mostly lead to non-synonymous changes (**Supplementary Table 3**). This could partly explain the strong bias towards non-synonymous (n=124) relative to synonymous (n=25) mutations seen in this line. Moreover, this line had a non-synonymous mutation in *recC*, which is involved in dsDNA repair and stress-induced mutagenesis (Al Mamun *et al.* 2012). Of note, a fluctuation test (Foster 2006) showed that the HMR and LMR lines have retained their high and low mutation rate, respectively, after the experiment (**Supplementary Table 4**).

An examination of the mutation pattern in coding regions with *dN/dS* ratio reveals no evidence for selection in both HMR and LMR lines (HMR lines: *dN/dS* = 1.014, site bootstrap 95%CI 0.962-1.065, LMR lines: *dN/dS*=1.093, site bootstrap 95%CI [0.812-1.354]), and a permutation-based test revealed no significant difference in *dN/dS* ratios between HMR and LMR lines (p=0.213). The use of PROVEAN scores (Choi and Chan 2015) to quantify the potential effects of amino acid substitutions and in-frame indels in coding sequences shows that a majority of non-synonymous (NS) mutations have a potentially strong effect on protein function for both HMR and LMR lines (61.3% and 64.8%, respectively, t-test, p=0.166, **Supplementary Fig. 3**). It suggests that the observed *dN/dS* ratio close to 1 in both HMR and LMR lines is not due to the observation of phenotypically neutral mutations, but that it is rather including many non-synonymous mutations that might impair cellular functions.

### Evolution of colony expansion speed

Theory predicts that an accumulation of deleterious mutations should lead to a reduction in expansion speed over time (Peischl *et al.* 2015). We indeed found that the colony size of HMR lines has significantly declined over time (-78 μm/day, 95% CI [-85; −70], p-value: <2 10^-16^), whereas the colony size of LMR lines has not significantly changed (-11 μm/day, 95%CI [-33; 10], p-value=0.29) (Fig. 3). Using the individual linear regression lines to estimate the relative change in colony size over the course of the experiment, we found that the colony size of HMR lines after 3 days of growth has decreased by 33±3% in 39 days (t-test, p-value<2.2 10^-16^). In contrast, colony size of LMR lines after the same period of growth did not significantly change (p-value=0.36), and the difference in colony size at day 39 is significant between the HMR and LMR lines (t-test, p-value=4.16 × 10^-7^).

### Reduced fitness of evolved strains

While the observed reduction in colony size over time is consistent with an accumulation of deleterious mutations due to random genetic drift (Peischl *et al.* 2015), an alternative explanation would be the establishment of adaptive pleiotropic mutations that increase the ability of bacteria to end up at the expansion front, while at the same time have a negative effect on growth rates. Such antagonistic pleiotropy between motility and growth related traits has recently been observed in experimental evolution studies of *Pseudomonas aeruginosa* (Van Ditmarsch *et al.* 2013). We therefore evaluated the fitness of HMR and LMR lines as their ability to grow competitively on agar. We let them directly compete with the ancestral strain during a radial expansion on an agar plate, and thus measured their relative fitness under the exact same conditions as in the 39-days experiment shown in Fig. 1 (**Supplementary Fig. 5**). We found that the ancestral strain clearly outcompeted all but one HMR lines (Fig. 4A), providing strong evidence for an overall decrease in fitness during the 39 days experimental evolution on agar. The LMR strains showed similar patterns, with an overall reduced ability to compete with the ancestral strain on agar (Fig. 4A). However, three out of ten LMR lines showed signals of adaptation to growth on agar as they clearly outcompete the ancestral strain (Fig. 4A). Importantly, these results demonstrate that adaptive process involving trade-offs between growth and motility are unlikely to be the main reason for the decrease in colony size over time. Rather, our results indicate that accumulated deleterious mutations strongly reduced the fitness of bacteria in HMR lines, and to a lesser extent in LMR lines.

### Alternative measures of fitness confirm predominant deleterious effects of mutations

To test whether mutations accumulated in the HMR lines had any negative impact on biological processes and thus on cellular functioning, we assessed the fitness of our evolved bacterial lines in two additional ways, complementing the radial competition assay (Fig. 4).

We first measured growth rate relative to a reference strain (different from the ancestral strain) during a linear expansion without any mixing (see **Supplementary Fig. 4A**). The advantage of this procedure relative to the radial growth competition described above is that one can directly transform differential growth rates into selection coefficients (Hosono *et al.* 1995; Korolev *et al.* 2012). One can thus more precisely assess the growth component of bacterial fitness than under radial expansion with mixed strains. However, note that with this measure of fitness, we ignore another component of fitness, which is the ability to make it to the wave front where bacteria can grow and reproduce (Korolev *et al.* 2012). In any case, we find that the fitness as measured by the mean relative growth rate of the HMR lines is significantly lower than that of the ancestral strain (*w̅* = 0.84, t-test, p=0.006, Fig. 4B). We note that 2 out of the 9 tested HMR lines show no reduction in fitness despite having accumulated many mutations (128 and 142 mutations). Interestingly, these two lines have non-synonymous mutations in the *mlc* (**m**akes **l**arge **c**olony) gene, known to enable prolonged growth on agar. Contrastingly, the average relative growth rate of LMR lines was slightly but not significantly larger than that of the ancestral strain (*w̅* =1.05, t-test, p=0.058), with 7/10 lines showing an increase in relative growth rate. Importantly, all lines with increased fitness (relative to the ancestral strain) in the competitive radial growth assay (Fig. 4A) also have an increased growth rate as measured by the linear competition on agar (**Supplementary Fig. 8**). This is again in contrast with a pleiotropic adaptation hypothesis for reduced colony size, under which we would expect to see lines showing a reduced growth rate measured by the linear growth experiment (Fig 4B) despite an increased overall fitness as measured by the radial competition experiments (Fig 4A).

We then estimated the fitness of bacteria as they grow in liquid batch culture (Fig. 4C). In that case, the fitness of HMR lines in liquid culture has significantly decreased relative to the ancestral strain (*w̅* = 0.859, t-test, p= 2.585e-15), whereas the fitness of the LMR lines has remained identical (*w̅* = 1.001, t-test, p=0.46).

### Distribution of fitness effects

We first estimated the distribution of fitness effects and the effective population size at the expansion front by fitting an analytically tractable model for the evolution of mean fitness during range expansions (Peischl *et al.* 2015) to the colony size trajectories shown in Fig. 3. The estimated DFE (Fig. 5A) is almost symmetrical around zero, with an average negative effect of 0.00379 per mutation. Furthermore, we estimate an effective population size at the expansion front of *N_e_* = 14.6, very close to previous estimates suggesting an effective size on the wave front of the order of 10 cells (Hallatschek *et al.* 2007; Korolev *et al.* 2011), and indicating that evolution at the expansion front is largely determined by genetic drift. Fig 5B illustrates simulations of the estimated model with examples of observed data and reveals a good fit between theory and data. Further, Fig. 5C shows that the difference between observed and expected fitness is not significantly greater than what would be expected by chance under our theoretical model of expansion load (p-value=0.18).

**Figure 5:**
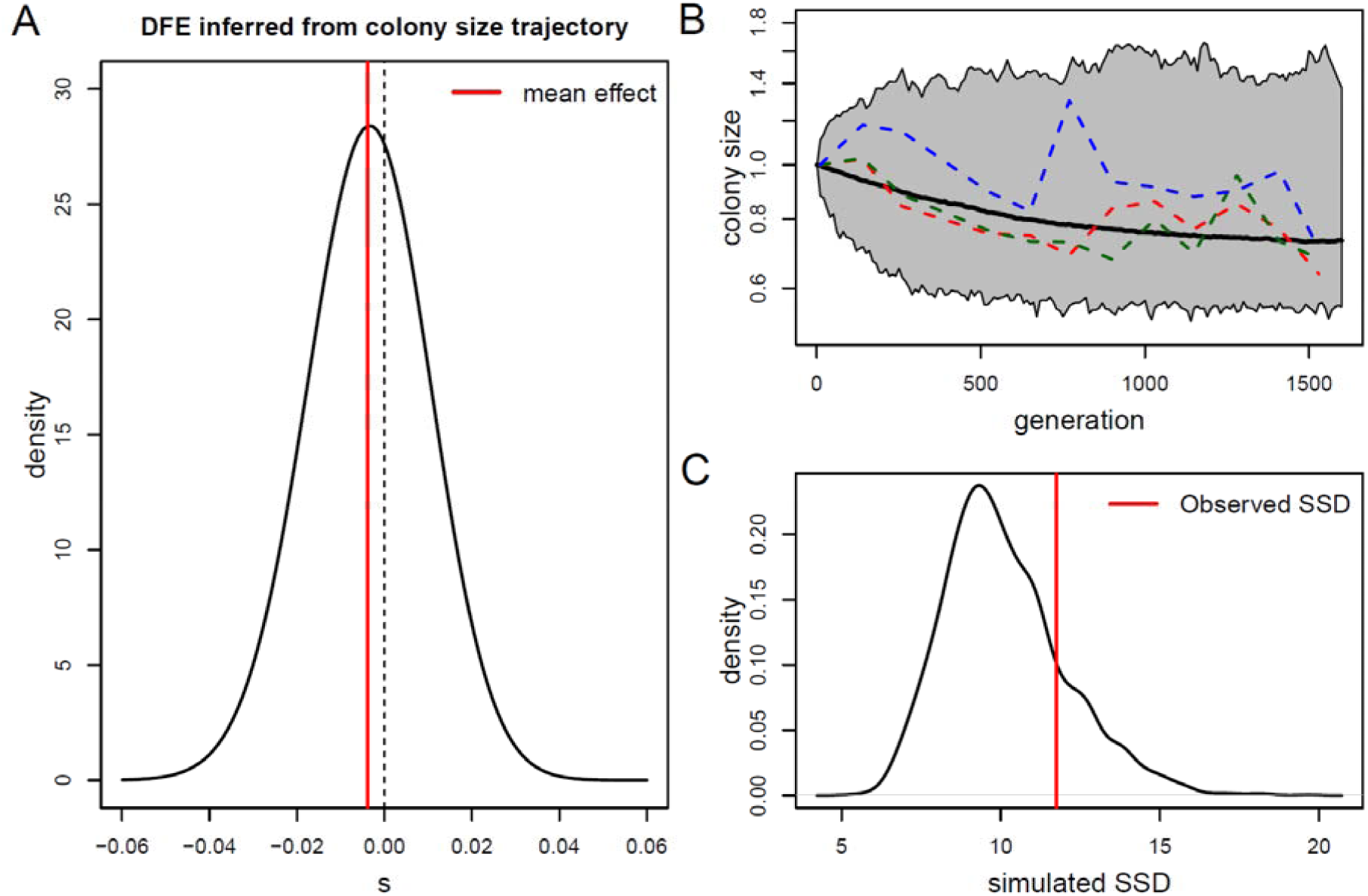
Distribution of fitness effects (DFE). **A**: DFE inferred from the evolutionary trajectory of colony size over time shown in Fig. 3. Parameters were estimated by minimizing the sum of squared deviations (SSD) from the expectation of colony size obtained from the model described in Peischl et al. (2015). Estimated parameters of the displaced Gamma distribution: *α* = 972; *β* = 2220.2; *δ* = 0.434. The effective population size at the expansion front is estimated as *N_e_* = 14.6, and the mean mutation effect is −0.00379. **B**: Evolution of colony size obtained by simulations using the estimated parameters. The solid line shows the average and the borders of the gray shaded area indicate the 0.5 and 99.5 percentiles of the simulated data, both estimated from 1,000 simulations. The dashed lines show 3 randomly chosen examples of the observed colony size evolution of the HMR strain. **C**: Test of goodness of fit of the observed fitness under the estimated DFE shown in pane A. The SSD between observed and expected fitness is compared to the distribution of SSD between the expected fitness and that simulated using the estimated parameters. The simulated SSD density was computed from 1,000 simulations. The observed deviation between expected and observed fitness in pane B is thus not significant (p-value=0.18).

The DFE estimated from the change in colony size over the 39 days of evolution as a function of the number of accumulated mutations is shown in Fig. 6A. This DFE is almost symmetrical around zero, and very similar to that shown in Fig. 5A, with an average negative effect of 0.0029 per mutation. This mean mutation effect is smaller than that shown in Fig. 5, since this DFE is only based on observed mutations, i.e. that were not lost due to selection during the time of the experiment, whereas the DFE reported in Fig. 5 considers all mutations, observed or not. Nevertheless, this DFE explains the data quite well, as shown on Fig. 6B, where we report the decline in fitness expected for this DFE and its 95% confidence interval under an additive model. Moreover, we see on Fig. 6C that the difference between observed and expected fitness is not significantly greater than what would be expected by chance under our model (p-value=0.80). We also tested whether the HMR and LMR lines have different DFEs using a likelihood ratio test, which reveals not significant (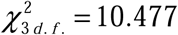, p-value=0.269), suggesting that the DFE in Fig. 6A can explain the dynamics of colony size change of both HMR and LMR lines.

**Figure 6:**
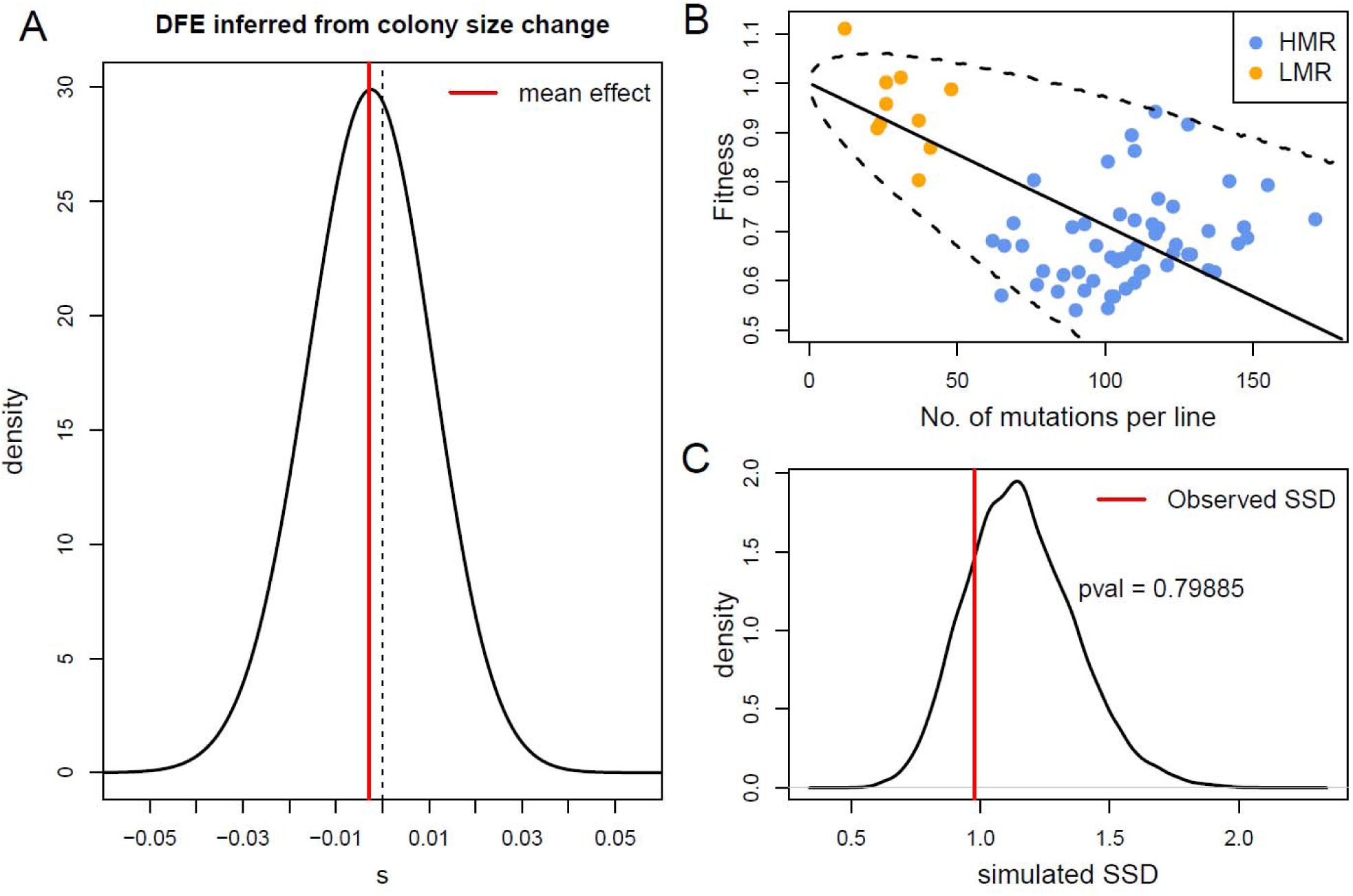
Distribution of fitness effects (DFE). **A**: DFE inferred from the change of colony size over time shown in Fig. 3. Maximum likelihood parameters of the displaced Gamma distribution: *α* = 989.95; *β* = 2357.2; *δ* = 0.417. Mean mutation effect = −0.00288. **B**: Fitness of bacteria as a function the number of observed mutations in HMR and LMR strains. The solid black line is the mean fitness decline expected under the DFE shown in pane A, and the dashed lines represent limits of a 95% confidence interval around the mean, both estimated from 50,000 simulations. **C**: Test of goodness of fit of the observed fitness under the maximum likelihood DFE shown in pane A. The sums of square deviations(SSD) between observed and expected fitness is compared to the distribution of SSD between the expected fitness and that simulated under the ML DFE for the same numbers of mutations as those observed. The simulated SSD density was computed from 20,000 simulations. The observed deviation between expected and observed fitness in pane B is thus not significant (p-value=0.80).

## Discussion

We provide here four lines of evidence that an accumulation of mutations during a range expansion leads to a decrease in fitness. We first observe that the ability to grow on agar of HMR lines significantly decreases over time in 52/57 lines, whereas that of LMR lines remain constant (Fig. 3). Second, direct competition of evolved strains with ancestral strains during a radial expansion that is like that occurring in our experimental setup reveals that 9/10 HMR lines and 7/10 LMR lines are outcompeted by the ancestral line (Fig. 4A). Third, a linear competition experiment, which allows us to directly estimate the fitness of the evolved strains relative to that of their ancestor, shows on average a significant 16% fitness reduction for HMR lines, and a non-significant increase in fitness for LMR lines (Fig. 4B). Finally, a measure of the growth ability of evolved strains in a completely different and nutrient-wise richer liquid medium shows a significant 14% reduced growth rate for HMR lines and again no change in fitness for LMR lines (Fig. 4C). Overall, these observations are in line with theory, which predicts that natural selection is relatively inefficient during range expansions due to the low effective size prevailing on the wave front, such that most deleterious mutations are not purged on the wave front (Peischl *et al.* 2013; Peischl *et al.* 2015). The use of this theoretical framework also allows us to directly infer the distribution of fitness effects from the reduction in growth rates of HMR lines over time under a hard selection model (Fig. 5A). The resulting DFE suggests that 60.3 % of mutations are deleterious, with a mean negative effect of 0.38% for new mutations. In addition, we have three lines of evidence suggesting that mutations have accumulated almost independently of their fitness effect. First, non-synonymous mutations have accumulated at the same rate as synonymous mutations, since *dN/dS* ratios are not significantly different from 1 for both HMR and LMR lines, and this even though most non-synonymous mutations are predicted to have a phenotypic effect using PROVEAN scores (**Supplementary Fig. 3**). Second, mutations have accumulated almost symmetrically along the genome on both sides of the origin of replication *oriC* (**Supplementary Figs. 6 and 7**), suggesting that the distribution of mutations along the bacterial genome reflects variable mutation rates that depend on positional constraints during chromosome replication (Foster *et al.* 2013) and not on local selective pressures. Third, the DFE inferred from the dynamics of colony size change as a function of the number of observed mutations per line is extremely similar to that inferred from all new mutations in HMR lines (compare Figs. 5A and 6A). This high similarity suggests that most new mutations have been retained during our evolutionary experiment, and that only highly deleterious mutations have been eliminated to lead to the 0.0009 shift between the estimated mean effects of new and observed mutations. The very low estimated effective size of ∼15 individuals on the wave front (Fig. 5) indeed suggests that only mutations with negative effects of the order of 1/15 (∼6.7%) or higher have been deterministically eliminated by selection.

The fact that the fitness of most HMR lines has decreased during their range expansion does not mean that all observed mutations are necessarily deleterious or neutral, as some mutations adaptive for the wave front conditions could have occurred. The estimated DFEs suggests that many positively selected and thus potentially adaptive mutations have indeed occurred (see Figs. 5A,6A). A closer examination of the early dynamics of the colony growths suggests that HMR and LMR lines could have adapted to life on wave fronts (Fig. 7). Indeed, using non-linear regression we observed that the colony size of LMR lines initially increases and then begins to decrease after 24 days of growth to reach levels similar to those observed at the onset of the experiment after 39 days of evolution (Fig. 7A). A similar analysis of the growth rates in the first 12 days of evolution of HMR lines shows that they initially slightly increase before declining after 6 days (Fig. 7B), in keeping with the potential occurrence of adaptive mutations in some HMR lines in the early phases of the experiment. For instance, some mutations could have potentially allowed some strains to preferentially occupy the wave front and thus access fresh nutrients. Such a positive selection of mutations allowing bacteria to occupy the expanding wave front have been recently described in several bacterial species (Oldewurtel *et al.* 2015; Van Ditmarsch *et al.* 2013). In a different setting, it has also been shown that beneficial mutations can very quickly increase in frequency during range expansions on agar, and this significantly faster than in well-mixed populations, so that beneficial mutants can rapidly colonize a large proportion of the colony wave front (Gralka *et al.* 2016a). However, by doing so, positively selected bacteria would enter an environment where local population size is low (estimated *N_e_* = 14.6, Fig. 5), genetic drift is high and selection is inefficient, making it more difficult to purge further deleterious mutations (Hallatschek and Nelson 2010; Peischl *et al.* 2013; Peischl *et al.* 2015). The evolution of *E. coli* strains during their growth on agar could thus result from a complex interplay between beneficial and deleterious mutations, but the effect of the later ones seems to predominate in most HMR lines after 39 days. The estimated overall 15% fitness disadvantage of HMR lines relative to the ancestral strain (Fig. 4B) makes them rapidly outcompeted by ancestral strains, and most of them entirely disappear from colonizing wave fronts during a radial competition experiment (Fig. 4A). This strongly indicates that population expansions on solid surfaces are detrimental for mutator strains, which could explain why they are not often seen in natural conditions (Matic *et al.* 1997; Tenaillon *et al.* 2000).

**Figure 7:**
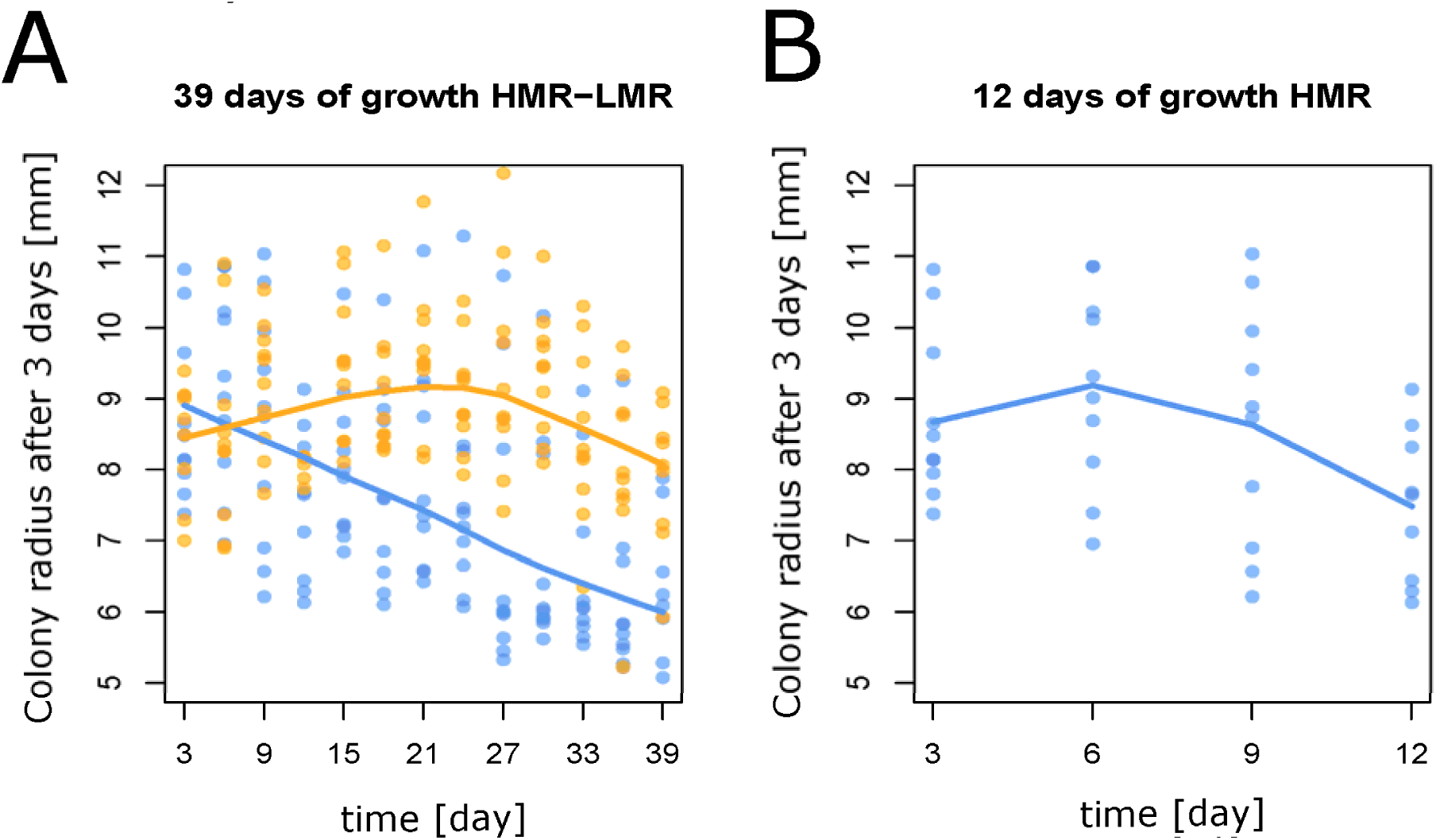
Early growth dynamics of HMR and LMR lines. **A** Evolution of HMR (experiment 2) and LMR colony size after 3 days of growth over the course of the experiment. The average size of HMR colonies estimated by a LOESS regression linearly declines over time, whereas that of LMR colonies increases until day 24, and then declines until the end of the experiment. **B**: Same as A, but only for HMR lines during the first 12 days of growth, showing a pattern similar to that of LMR lines but on an approximately 4 times shorter time scale, which approximately corresponds to their ∼3.7 times higher mutation rate (see **Supplementary Table 4**).

Since HMR lines show reduced growth abilities not only on agar but also in a well-mixed liquid culture (Fig. 4C), some of the accumulated mutations are likely to be unconditionally deleterious. Thus, even though bacterial populations are generally considered to be able to adapt to almost any environmental conditions (Hindre *et al.* 2012), HMR strains seem to have accumulated deleterious mutations in a wide range of cellular processes. Our results thus imply that many mutations seen in natural bacterial populations are not necessarily adaptive and that populations of bacteria growing on two dimensional surfaces can develop an expansion load (Peischl *et al.* 2015), even though the speed of this process should be slower for non mutator strains. Our results thus highlight the importance of considering the spatially explicit process of bacterial growth when studying bacterial adaptation and evolution. The expansion load we demonstrate here in bacteria could also happen during the expansion of other populations, including humans (Henn *et al.* 2015), but it could also affect other types of expansions, like the growth of solid tissues in eukaryotes. The analogy between the evolution of bacterial communities and the growth of eukaryotic tissue has recently been highlighted (Lambert *et al.* 2011). Indeed, multicellular organisms go through millions to trillions of cell divisions during their life span, accumulating somatic mutations at a rate about ten times higher than germ line mutations (Lynch 2010; Shendure and Akey 2015), potentially contributing to cancers and other human diseases (Shendure and Akey 2015). In addition to having triggered the development of specific life-history traits in most organism (reviewed in Charlesworth and Charlesworth 1998), deleterious mutations could have also led to the development of specific cellular mechanism preventing their specific accumulation during tissue growth (e.g. apoptosis), which should be the object of further studies.

## Acknowledgments

We are grateful to Oskar Hallatschek, Mark Kirkpatrick, Sally Otto and Olin Silander for constructive comments and critical reading of an earlier version of the manuscript and to Jean-Frangois Flot for discussions at various stages of this project. We also want to thank Tosso Leeb, Cord Drögemüller and the NGS core facility of the University of Berne for their support. LB, ID, and SP were partly supported by Swiss NSF Grants No. 31003A-143393 and 310030B-166605 to LE and MA.

